# The Impact of Coiled-Coil Domains on the Phase Behavior of Biomolecular Condensates

**DOI:** 10.1101/2024.12.12.628163

**Authors:** Zhouyi He, Jens-Uwe Sommer, Tyler S. Harmon

## Abstract

Spatial organization is fundamental to biological systems, with biomolecular condensates as a key subset. Many studies show that folded domains play important roles in condensate formation by facilitating interactions. However, little is known about how the presence of large structured elements impacts condensate formation. Using coarse-grained simulations, we investigated a model system of two multivalent proteins, one containing a coiled-coil domain (CCD), which undergoes liquid-liquid phase separation (LLPS). We found that CCDs promote LLPS by preventing loop-closure defects, enabling protein networking. Replacing the CCD with a flexible linker abolishes LLPS due to formation of oligomeric clusters. There is a critical length of the CCD where the system rapidly changes from no LLPS to LLPS at low concentrations. This highlights their potential in regulating condensate formation and properties. This study provides insights into the phase behavior of biomolecular condensates and offers a framework for designing synthetic condensates with tunable phase behaviors.

---

Organelles are specialized sub-units within cells that perform specific functions. Traditionally, they are viewed as membrane-bound structures enclosed by lipid bilayers. However, recent research has highlighted the significance of non-membrane-bound organelles, also known as biomolecular condensates (condensates). ^1–7^ They form through liquid-liquid phase separation (LLPS)[simulation papers], providing a fundamental mechanism for cellular organization and playing crucial roles in various cellular processes.

Many studies have focused on how flexible intrinsically disordered regions drive LLPS. ^8–11^ In contrast, coiled-coil domains (CCDs), a particular type of structured motif, are rigid and are found in many phase-separating proteins. ^12^ Coiled-coils are bundles of alpha-helices that are coiled together like the strands of a rope, forming rod-like structures.^13–19^ The significance of CCDs in the literature is primarily associated with their role as spacers, such as in EEA1 tethers or as components of motor proteins.^20–22^ However, the role of CCDs in condensate formation has received little attention.

Here, we show that CCDs promote LLPS and condensate formation. Our work is motivated by a recent experimental study by Heidenreich *et al*., where they designed CCD-containing proteins that phase separate into synthetic condensates in yeast. ^23^ This system consists of two components: One protein contains a homo-tetramerization domain and a hetero-interacting domain (E9), which primarily exists as a tetramer. And the other contains a homo-dimerization domain and a hetero-interacting domain (Im2), which primarily exists as a dimer (Figure 1a). Notably, the homo-dimerization domain is a CCD. ^13^ These proteins interact through the hetero-interacting domains (E9:Im2), forming a condensate (Figure 1d).

**Figure 1:**
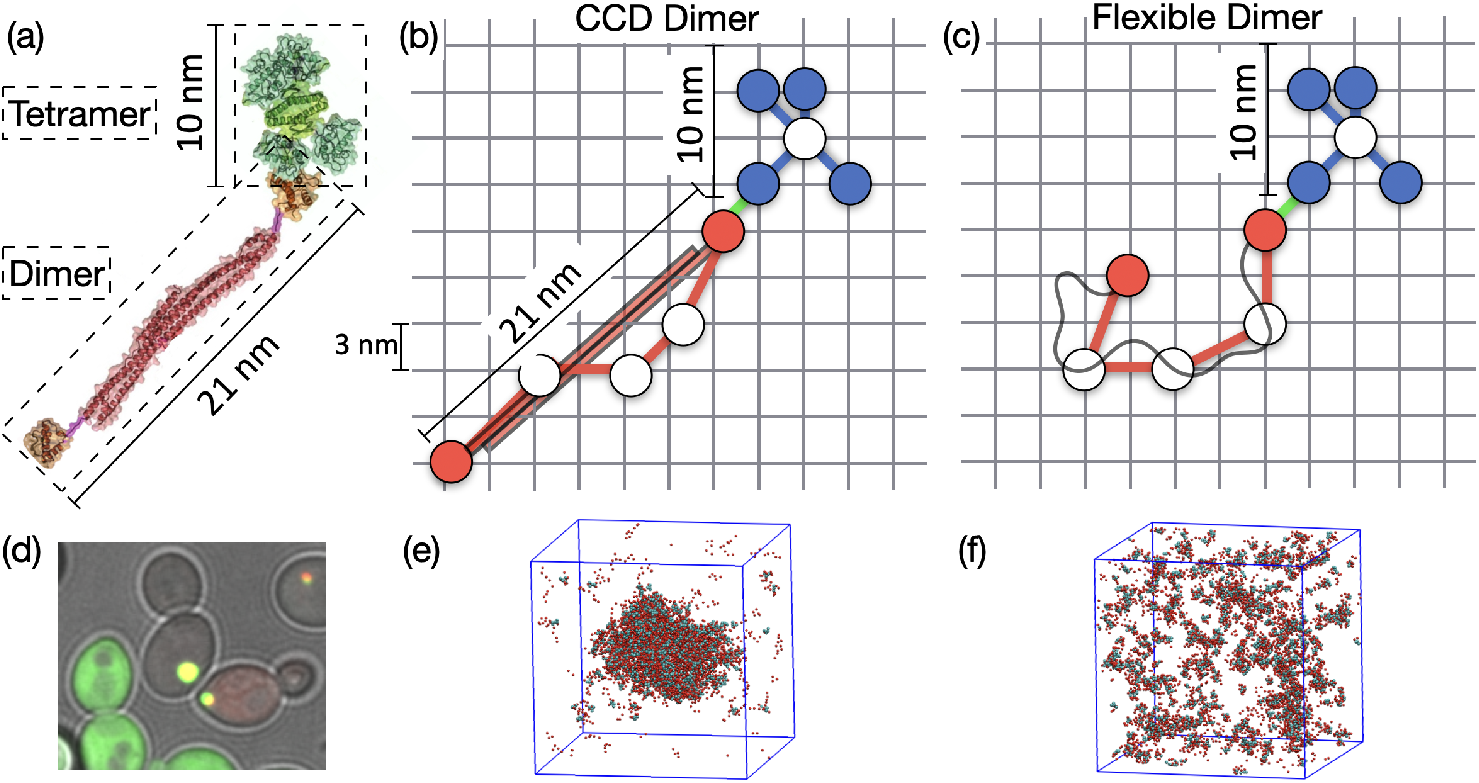
Tetramer-Dimer Model System. (a) Crystal structures of the proteins designed by Heidenreich et al.^23^ Tetramers containing tetramerization domains (green, PDB: 1AIE) and E9 domains (cyan, PDB: 2WPT). Dimers containing CCD dimerization domains (red, PDB: 4LTB) and Im2 domains (orange, PDB: 2WPT).^23^ (b) and (c) Coarse-graining protein oligomers onto a lattice. The tetramer is modeled as a star polymer with five beads, while the dimer is modeled in two ways: (b) as a CCD Dimer with a harmonic elongation potential applied to maintain its extended structure, and (c) as a Flexible Dimer without elongation restraints. Each lattice site measures 3 nm. (d) Condensate formation observed in yeast. Bright yellow circles inside the cells are condensates of designed proteins; Cells that are completely green are only expressing tetramers; Dark cells are expressing neither. For more details on the experimental methodology, refer to Heidenreich et al. ^23^ (e) and (f) Simulation snapshots of 3D lattice simulations of the coarse-grained oligomers. (e) The CCD Dimer system shows clear LLPS, (f) the Flexible Dimer system does not phase separate.

This system can be viewed in analogy to the formation of polymer gels, which consist of a star polymer with 4-functional branches and a linear polymer with 2 functional ends.^24^ In contrast to gel formation, which generally occurs through percolation,^25^ the condensate formation is LLPS. As shown below, the elongation of the CCD is crucial for this process. Without the CCD, the system forms stable oligomeric clusters at low concentrations and gelates at high concentrations (Figure 2b). In contrast, the long CCD prevents the formation of stable small clusters due to geometric constraints, leading to LLPS (Figure 2a). We employed 3D lattice polymer Monte Carlo simulations,^26,27^ where protein oligomers were coarse-grained and represented by a bead-spring model with excluded volume. To represent the dimensions of folded domains in real systems, ^28,29^ the lattice sites are associated with a length scale of approximately 3 nm. This level of coarse-graining allowed us to simulate systems with a length scale as large as 1 *µ*m with 10,000 polymers. Interactions between beads are modeled by formation of reversible bonds corresponding to protein domain:domain interactions. The Monte Carlo moves in our simulations include: 1) local-scale moves where one or two beads move in space and change interaction partners, 2) polymer-scale moves where a polymer slithers or moves translationally, and 3) global-scale moves where clusters of polymers are moved as a group to anywhere else in the simulation. For more detailed descriptions see supplementary information (SI).

**Figure 2:**
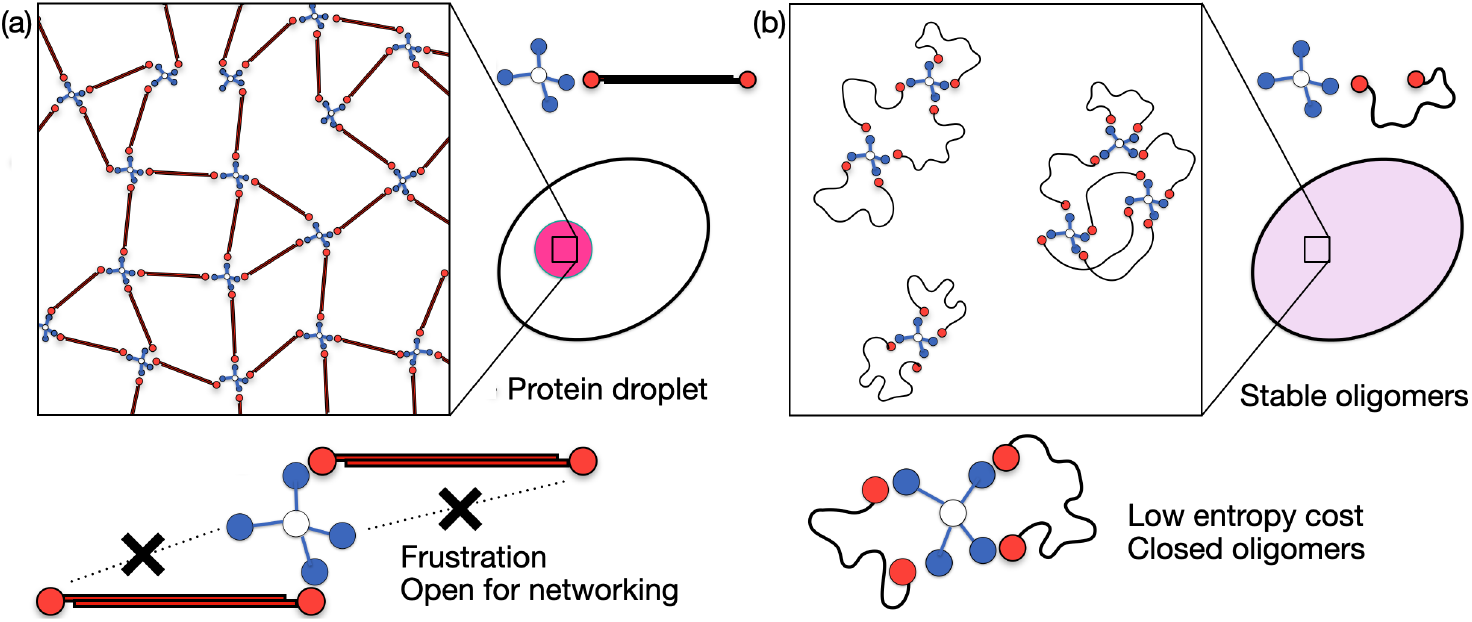
Coiled-Coil Domains (CCDs) Promote Phase Separation through Geometric Constraints. (a) Cartoons of CCD Dimers interacting. The presence of CCDs creates geometric frustration, preventing a tetramer from closing with two dimers. This configuration allows each dimer to connect multiple tetramers, facilitating network formation coupled with LLPS. (b) Cartoons of Flexible Dimers interacting. Tetramers can easily be closed with two dimers that do not contain CCDs. This leads to small oligomeric clusters, limiting network formation and preventing LLPS.

The tetramer is coarse-grained as a 4-branch star polymer of 5 beads with a total size of 10 nm. The center bead represents the tetramerization domains and the four branching beads are the E9 domains (Figure 1b, c). The dimer is coarse-grained as a linear polymer of 5 beads with a contour length of 21 nm. These beads represent the CCD with the end beads also including the E9 domains (Figure 1b, c). These choices ensure that the coarse-graining reflects the real length scales of the protein oligomers.

To capture the rod-like geometric constraint introduced by the CCD, we added an elongation potential to constrain the distance between the two end beads of the dimer: *U* = *K*(*L − L*_*eq*_)^2^, where *L* is the distance, *L*_*eq*_ is the equilibrium distance, and *K* is the elongation potential coefficient. We note two important limiting cases: 1) ‘CCD Dimers’ where *L*_*eq*_ = 21 nm, the system reproduces the extension of the CCD (Figure 1b), and 2) ‘Flexible Dimer’ when no elongation potential is applied, the system behaves as if the CCDs are replaced with flexible linkers of the same contour length (Figure 1c). For a more detailed description of the simulation designs, parameters, and simulation convergence see the SI.

Large scale simulations of mixing tetramers with CCD Dimers behave qualitatively different compared to mixing with Flexible Dimers. Tetramers with CCD Dimers form large condensates consistent with experimental observations (Figure 1d and e). On the other hand, tetramers with Flexible Dimers do not form condensates (Figure 1f).

These results can be qualitatively understood by considering the geometrically allowed microscopic arrangements of the polymers. For the Flexible Dimers, the average end-to-end distance is 13 nm, which is about the size of the tetramers. Therefore, the two branches of one tetramer could be closed by one dimer forming a loop defect, which accounted for around 25% of all tetramer interaction sites (Figure SI 4). Hence, small clusters are prevalent in dilute conditions, preventing further expanded networks associated with LLPS (Figure 2 b).

In contrast, with CCDs, the intramolecular interactions are restricted because the length of the CCD separating the ends of the dimer (21 nm) exceeds the length of the tetramer (10 nm). The closure of tetramers by one dimer is now geometrically forbidden. This promotes the formation of protein networks and LLPS into condensates (Figure 2a). At higher concentrations, a gelation transition is expected in both cases. For further discussion about the microscopic arrangements see SI.

To quantitatively understand the impact of the CCDs we used simulations with long rectangular periodic boxes (0.15 × 0.15 × 1.2*µm*^3^). Condensates will form a slab spanning the shorter dimensions (x,y) while phase separation takes place in the longer dimension (z). This reduces the impact from surface tension and simplifies extracting concentrations of coexisting phases (Figure 3a and SI).

**Figure 3:**
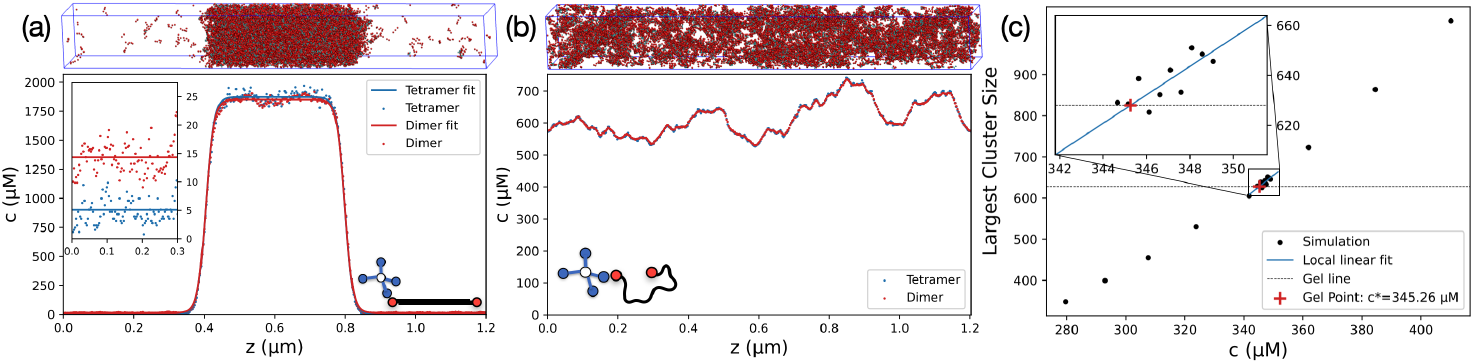
Quantifying Phase Separation and Gelation. (a) Snapshot of a SLAB simulation for CCD Dimers and the average concentration profile along the z-axis. This system undergoes LLPS, forming a dense phase with a concentration of 1850 *µ*M and a dilute phase with a saturation concentration *c*_*Sat*_ of 10 *µ*M. Note the center of mass of each frame is shifted to the center of the box before averaging in order to align the condensate phase. (b) Snapshot for Flexible Dimers and the average concentration profile along the z-axis. This system only forms a gel at high concentrations without undergoing LLPS. Simulations in (a) and (b) have average concentrations of [*Tetramer*] = [*Dimer*] = 615 *µ*M. (c) Quantification of the gel point. Each point corresponds to a simulation at a different average concentration. The gel point is calculated by interpolating the cluster size to the cutoff cluster size (dotted line *≈*630) that is motivated by Flory-Stockmayer theory, ^3,27,30–32^ see SI.

Some systems will not phase separate but instead undergo a gelation transition at high concentrations (Figure 3b). We use a cutoff for the fraction of components in the largest connected cluster that is motivated by Flory-Stockmayer theory^30–32^ to define systems that are gelated (Figure 3c and SI).^27,33^ If the system is a gel and has a uniform concentration across the simulation box then it has undergone a gelation transition without LLPS but through the pathway of percolation(Figure 3b and SI). Consistent with the previous results, the Flexible Dimers could only form a gel spanning the whole system.

Simulating tetramers with the CCD Dimers shows robust LLPS with concentrations around 200 times higher in the condensate phase compared to the bulk (Figure 3a).

Heidenreich et al. experimentally measured the two-dimensional saturation concentrations *c*_*Sat*_ (binodal) for the synthetic condensates in yeast (orange line in Figure 4a). We calculated the phase diagram by simulating different average concentrations of two components for both limiting models (Figure 4b and c). Additionally, we found where the system gelates outside the binodal.

**Figure 4:**
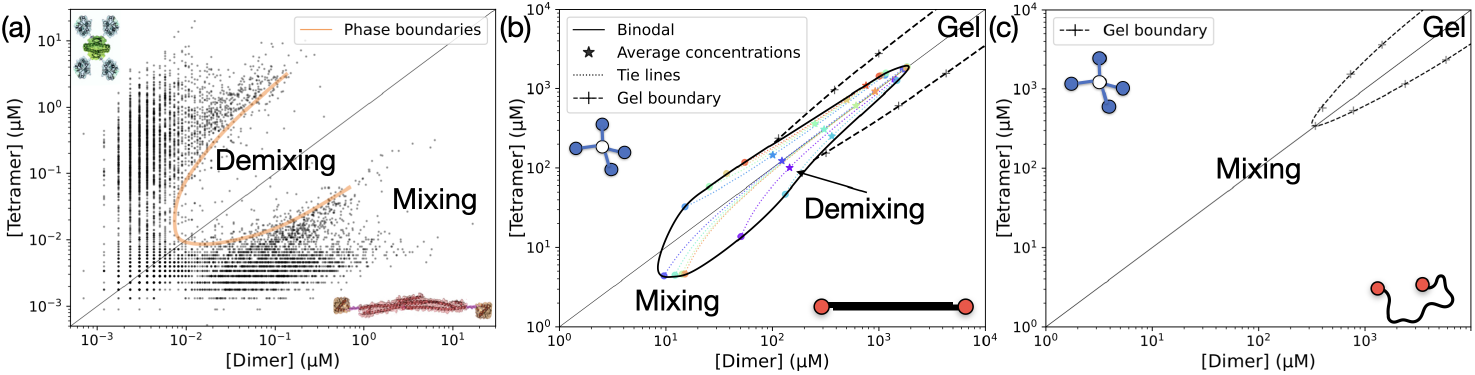
Comparing Phase Diagrams. (a) Dilute phase concentrations for tetramers and dimers measured from 6818 cells (black dots, data from Heidenreich et al.^23^). The experimental saturation concentrations are highlighted in orange. (b) Phase diagram from lattice simulations for CCD Dimers at equilibrium length *L*_*eq*_ = 21 nm, corresponding to the designed CCDs. Simulations with average concentrations in the demixing region robustly phase separate. Each average concentration and its corresponding dense and dilute phase points are connected by a color-matched tie line (dotted line). The binodal (phase boundary) is interpolated from the resulting concentrations of the two coexisting phases in the demixing region. Simulations in the gel region form a gel without LLPS. The gel lines were calculated for multiple concentration ratios between tetramers and dimers, see Figure SI 6a and b. The gel lines connect to the phase diagram’s plait points (where the dilute and dense concentrations meet). (c) Phase diagram for the Flexible Dimers. These simulations only exhibit a gel transition without demixing, see Figure SI 6c and d. The diagonal lines indicates the 1:1 stoichiometry.

The resulting CCD Dimer phase diagram qualitatively agrees with the experimental data (Figure 4b). We used weaker interaction strengths than the experimental system to calculate the binodal curve and gel lines for computational reasons, see SI.

Results with 1:1 stoichiometry with stronger interactions are shown in Figure 6a and SI, which show that the simulation saturation concentrations are one order of magnitude higher than the experimental results. This can be explained by factors outside the scope of our minimal model. First of all, most proteins in cells involved in condensates display weak non-specific attractive interactions. Thus the dense (condensate) phase is formed at a lower saturation concentration. Similarly, the cytoplasm is a crowded environment with other biopolymers, which increases the effective chemical potential, thereby lowering the saturation concentration compared to our simulations. Finally, the dissociation constants of E9:Im2 are measured in vitro which are different solution conditions from the cellular environment. We note that, in particular, non-specific interactions can be implemented into the model in further studies.

In contrast to the CCD Dimers, Flexible Dimers lead to gel formation but never to LLPS, which is characterized by the gel line in Figure 4c. In both cases LLPS/gelation is most salient at 1:1 stoichiometry, as expected.

We further investigated how the length of the CCD in the dimer impacts phase behavior by varying *L*_*eq*_ from 3 nm to 33 nm (Figure 5a inset). We found that the phase behavior changes rapidly around a critical dimer length 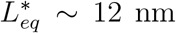 nm (Figure 5b). Systems with dimers longer than this phase separated robustly. Systems with dimers shorter than this did not phase separate and only gelated.

**Figure 5:**
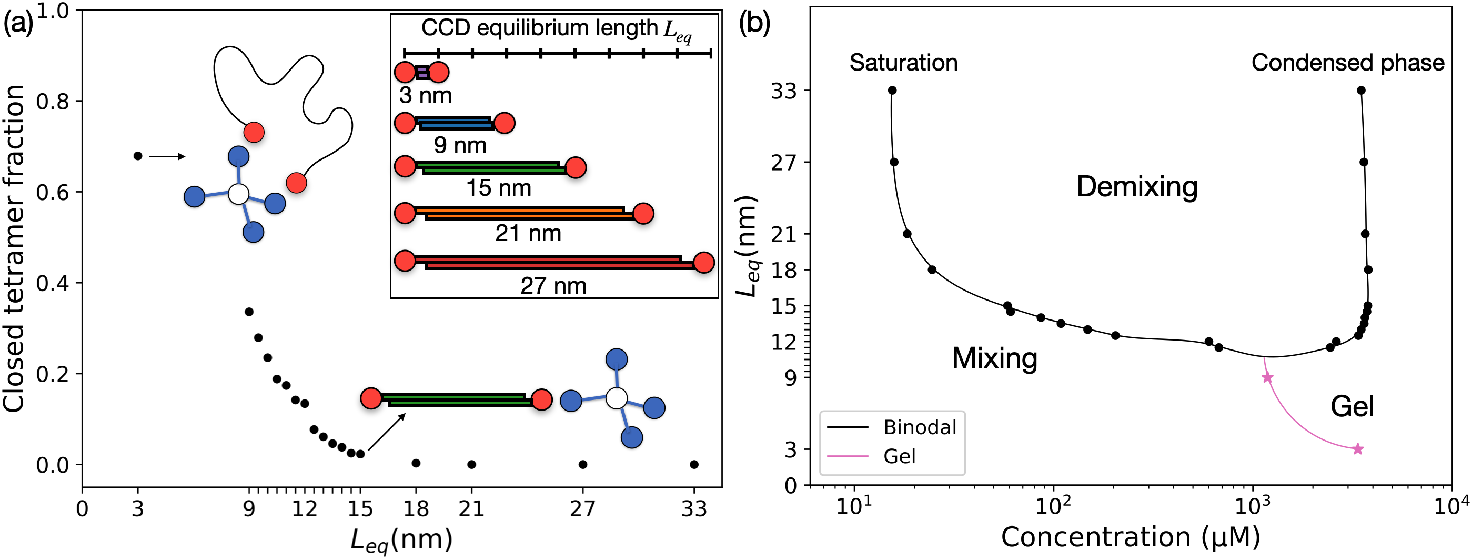
Influence of CCD lengths on Phase Behavior. (a) Fraction of tetramer interaction sites in a closed state (loop defect) as a function of equilibrium lengths *L*_*eq*_. We define a closed state as when the dimer forms a loop with the same tetramer and hence cannot contribute to the connectivity of the condensed state.^24^ Longer CCDs reduce self-loop closure of tetramers. Upper right: Schematic of dimers with different CCD lengths, *L*_*eq*_. Phase diagram as a function of *L*_*eq*_ at 1:1 stoichiometry. The saturation concentration changes by over an order of magnitude from *L*_*eq*_ of 12 nm to 18 nm. The system can only form a gel when *L*_*eq*_ is less than 12 nm (magenta line). (c) All simulations use an elongation potential coefficient 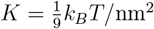; for results at other values of *K* see SI.

This critical length scale emerges because it matches the length between the interaction domains of the tetramer *∼* 10 nm. Therefore, when the CCD Dimer is longer than 12 nm, bond formation is geometrically frustrated as one tetramer cannot interact with both ends of one dimer (Figure 5a). Such loop defects are known to be important for shifting the gel point in polymer networks formed by flexible star polymers.^24^

As *L*_*eq*_ increases from 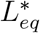, the saturation concentrations shift to lower concentrations by nearly two orders of magnitude (Figure 5b). Thus, variation of *L*_*eq*_ is an effective way to control the phase behavior of systems with CCDs. Additionally, *L*_*eq*_ sets the density of the resulting condensates, which can be used as a feature to control the molecular transport dynamics and add selectivity over molecules of certain sizes.^34^

Condensate formation is dependent on the the E9:Im2 binding strength, characterized by its dissociation constant *K*_*d*_. For intermediate binding strengths, the system phase separates with a linear relationship between the *K*_*d*_ and the *c*_*Sat*_. This linear behavior corresponds to an exponential change in *c*_*Sat*_ as a function of binding energy, see SI. A minimum binding strength, corresponding to a maximum *K*_*d*_, is required for the system to phase separate, beyond which the system will only form a gel at high concentrations (Figure 6a).

**Figure 6:**
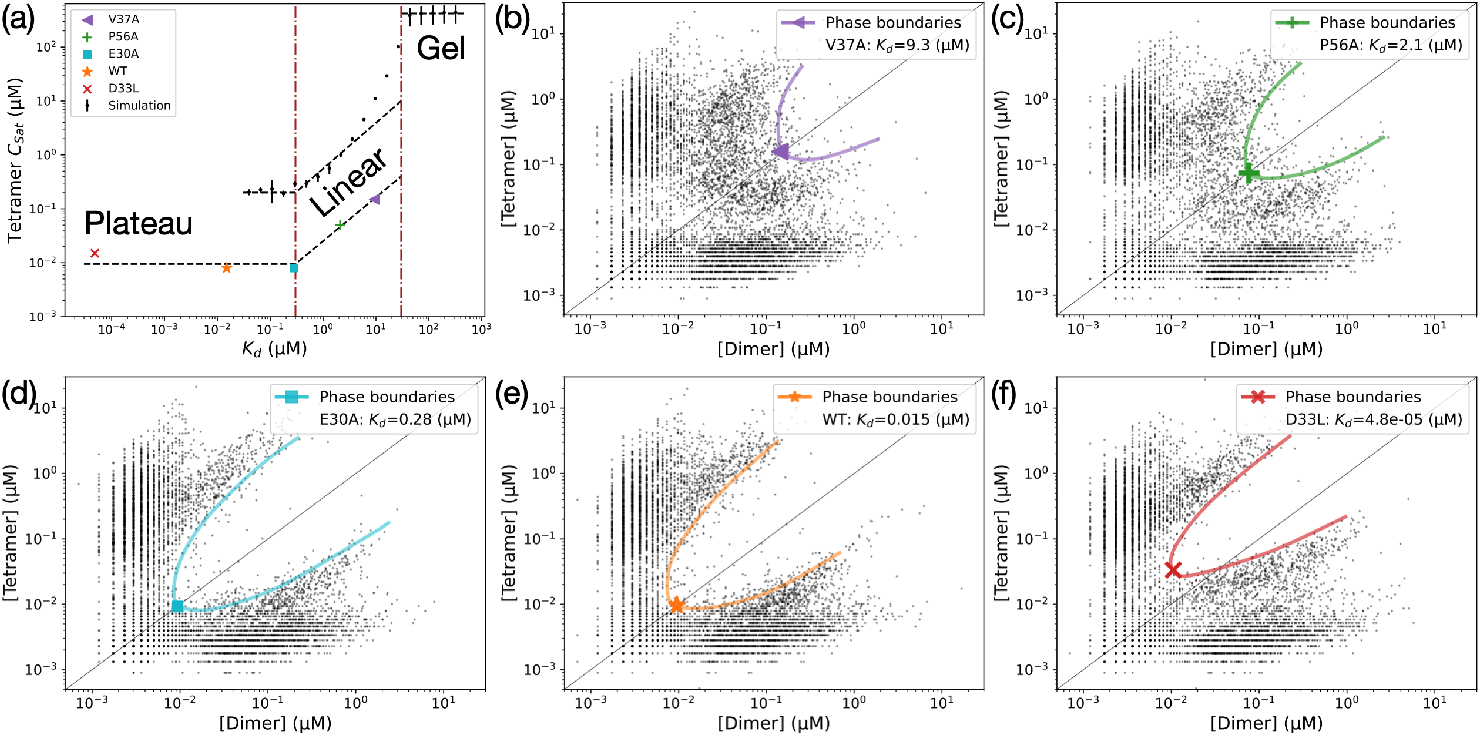
Phase Behavior for Different Binding Strengths. (a) Saturation concentrations of tetramer, *c*_*Sat*_, at 1:1 stoichiometry as a function of the E9:Im2 dissociation constant, *K*_*d*_. See SI for the conversion from simulation binding energies to *K*_*d*_ values. Simulations are shown in black where the error bars are the standard deviation from 6 simulations. There are three distinct regions: Plateau, Linear, and Gel, indicating different phase behaviors as binding strength changes (dashed lines are a guide to the eye). Experimentally derived *c*_*Sat*_ values for the wild type (WT) and four E9-Im2 variants (V37A, P56A, E30A, and D33L) are annotated in color for comparison.^23^ (b-f) Dilute phase concentrations for tetramers and dimers measured from *∼* 6500 cells for each variant with different *K*_*d*_ values (black dots, data from Heidenreich et al.^23^). The experimental saturation concentrations are highlighted in color. The symbols show the lowest saturation concentrations and are plotted in (a).

Moreover, there is also a maximum binding strength that impacts *c*_*Sat*_. The value of *c*_*Sat*_ plateaus for any stronger interactions (lower *K*_*d*_). This can be explained as follows: Strong binding strengths lead to very low saturation concentrations. While geometric constraints from elongated CCD Dimers prevents loop defect formation, at very low concentrations oligomeric clusters of higher order start to dominate which satisfy interactions without net-working (Figure 2b). Thus, our simulations reveal that formation of oligomeric clusters sets a absolute lower bound for *c*_*Sat*_ with a specified molecular architecture.

Our results are confirmed by the experiments by Heidenreich et al. where phase diagrams with different affinities are reported.^23^ The corresponding *K*_*d*_ ranges from 10^*−*5^ *µ*M to 10 *µ*M^35^ (Figure 6b-f). Their results fit into two of our three regimes - linear and plateau (Figure 6a). With zero parametrization, the binding strength for the transitions between linear and plateau are identical between the simulations and experiments. Based on our study, we expect that experiments with weaker E9:Im2 interactions would push system into the gel regime.

In this study, we investigated the phase behavior of a system composed of a dimer and a tetramer component. In particular we have focused on the impact of coiled-coil domains (CCDs). We found that systems with Flexible Dimers could not undergo LLPS but only gelation, whereas CCD Dimers robustly phase-separated. We can explain this behavior by the frustration of defect structures: a tetramer cannot interact with both ends of a long rigid dimer, thus cannot form loop defects. This leaves interaction sites open for further network formation, which drives LLPS.

In contrast, with flexible dimers, a tetramer can interact with both ends of a dimer, forming a loop defect with fully satisfied binding sites at concentrations below the overlap threshold. Our analysis further predicts a minimal saturation concentration as binding strength is increased. In this case, higher-order loop defects dominate, seen as two tetramers and four dimers forming a stable cluster. This transition can be observed in both simulations and experiments. The suppression of LLPS for the Flexible Dimer occurs because one scaffold protein’s valency is an integer multiple of the other’s (4:2), which is an example of the so-called ‘magic-number effect’.^36^ However, we also demonstrate that the magic-number effect is not intrinsic but only valid if both the valence and geometry are compatible. Sufficiently flexible linker elements are a way to guarantee geometric compatibility, as seen in the Flexible Dimers.

CCDs are abundant in RNA-binding proteins present in biomolecular condensates and could have important role for the formation and function of condensates. Additionally, post-translational modifications are used to control the formation/stiffness of CCDs ^37^ and thus could regulate the condensate formation or density.

These insights also provide a framework for predicting and manipulating gelation and LLPS in synthetic multi-component systems. We show that the phase diagram of this system could be directly tuned by introducing CCDs of different lengths. Going beyond the present experimental studies, our phase diagram includes the dense branch of the binodal, the density and composition of the condensate, which is important for functionalizing these condensates. The density is connected to the viscosity and recruitment of different client molecules. The condensate composition defines the local chemical environment which impacts recruitment specificity and chemical reactions. This could pave the way for designing synthetic biomolecular condensates for applications in biotechnology and medicine.

This work highlights the need for mapping the general principles of how structural domains contribute to LLPS beyond acting as interaction sites.

## Acknowledgement

We thank Emmanuel Levy and Meta Heidenreich for providing experimental raw data used for Figure 1d, 4a and 6b-f and insightful discussions. We are grateful to Simon Alberti, Titus Franzmann, Theo Wallenfang for collaboration in a related project. We appreciate Patrick McCall, Xi Chen, Arash Nikoubashman, Mrityunjoy Kar, Michael Lang, Holger Merlitz, and Ron Dockhorn for useful discussions. The authors acknowledge the Center for High-Performance Computing (ZIH) Dresden for granting simulation time.

