## Supporting_Information for "The Impact of Coiled-Coil Domains on the Phase Behavior of Biomolecular Condensates"

**Contents:**

- Lattice simulation methodology
- Convergence of simulation
- Determining gelation transition and liquid-liquid phase separation
- Impact of elongation potential on phase behavior
- Conversion of binding energy to $K_d$

### Lattice simulation methodology

#### Algorithm details

The code of the Monte Carlo lattice simulation engine for polymers used in this work is developed internally and is available with a user guide at GITHUB Repository.[1] It models the polymers as a set of beads connected with linkers on a lattice. The beads move around and interact with beads on adjacent lattice sites. For typical phase-separating systems there will be a single large cluster (condensate) at equilibrium.

The interactions for the beads are modeled as 'transient bonds' with neighboring beads. Beads are either bonded to one other bead or unbonded. Beads can never be bonded to more than one bead at any given time. The strength of such pairwise bond is characterized by the bond energy.

For each step one of seven moves are attempted. They were designed based on what we thought would get the system to equilibrium as fast as possible without consideration for realistic dynamics.

- **Rotation move**

  Randomly choose one bead. If it has a bond with another bead, break this bond. Randomly form a new bond with an unbonded neighboring bead or remain unbonded.

- **Translation move I**

  Randomly choose one bead. If it has a bond with another bead, break this bond. Translate this bead to another position, within geometrical constraints. Randomly form a new bond with an unbonded neighboring bead or remain unbonded.

- **Translation move II**

  Randomly choose one bead.

  -If it has a bond with another bead, translate this pair to another position, within both of their geometrical constraints.

-If it is unbonded, translate this bead to another position, within geometrical constraints.

- **Slither move I**

  This is only used for homopolymers. Randomly choose one polymer. Remove one bead from one end of the polymer and add a bead to the other end. For the new bead, randomly form a new bond with an unbonded neighboring bead or remain unbonded. (Equivalent to the polymer slithering 1 bead.)

- **Slither move II**

  This is only used for linear polymers and polymers with the identical linkers. Randomly choose one polymer. Remove $x$ ($x < 5$) beads from one end and add $x$ beads to the other end. Break all the bonds the polymer has. For each bead in the polymer, randomly form a new bond with an unbonded neighboring bead or remain unbonded. (Equivalent to the polymer slithering $x$ beads.)

- **Cluster move I**

  Randomly choose one cluster, defined as all polymers that are connected through bonds. Translate the cluster to another position anywhere in the lattice.

- **Cluster move II**

  Randomly choose one polymer. Calculate the cluster that it is in. If the cluster has fewer than 6 polymers, translate the cluster to another position anywhere in the lattice.

#### Simulation details

We used the SLAB technique, where the z-axis of the simulation is much longer than the other dimensions (50*50*400 lattice sites (LS)). We ensured that there was enough material to form a condensate that spanned the (x,y) dimensions if LLPS occurred. If the condensate grows, the slab gets thicker along z-axis. This technique reduces the impact of surface tension on the condensates properties, because the area of the condensate and is invariant to the volume of the condensate.

We simulated 2500 tetramers and 5000 dimers, unless otherwise noted. The system consists of $2500 \times 5 + 5000 \times 5 = 37500$ beads. The concentrations reported are in terms of binding site concentration, to highlight the 1:1 stoichiometry ($[2500 \times 4 : 5000 \times 2] = [10000 : 10000]$).

We used a binding energy between the E9 and Im2 beads of $6k_BT$, unless otherwise noted. A system typically converges within $10^{10}$ steps. Therefore, our simulation steps are typically at the order of $10^{11}$.

There are two ways of setting up the initial conditions for the simulation. 1) Start dilute: all polymers are randomly distributed in the box forming a homogeneous phase. 2) Start dense: all polymers are randomly distributed within a small height in the simulation box forming a dense phase.

The elongation potential constrains the distance between the two end beads of the dimer: $U = K(L - L_{eq})^2$, where $L$ is the distance, $L_{eq}$ is the equilibrium distance, and $K$ is the elongation potential coefficient. We note two important limiting cases: (i)'CCD Dimers' setting $L_{eq} = 21$ nm, the system reproduces the extension of the CCD (Figure 1c), and (ii)'Flexible Dimer' when no elongation potential is applied, the system behaves like a system where the CCDs are replaced with flexible linkers of same contour length.

The elongation potential coefficient $K$ for all simulations in main text is set as $1k_BT/LS^2$. For the impact of $K$ on phase behavior, see later section.

### Convergence of simulation

We demonstrate the robustness of our simulation results by investigating the convergence of the simulations under two different starting conditions, "dilute" and "dense" described above. We show the equilibration time is typically within 30 percent of the simulation trajectories (Figure SI 1 and 2). All reported equilibrium data is averaged over the later half of the simulation.

Figure SI 1 shows the energy and the largest cluster size converge within 10 percent of the simulation for both initial conditions and both types of polymers.

When analyzing concentration profiles of LLPS, for each frame, we shifted all the z coordinates such that the center of mass of the largest cluster $z_{Cluster}^{CoM}$ is at the center of the simulation box $L/2$ (re-center).

$$z_{Cluster}^{CoM} = \quad \mathrm{mod}\left(\frac{\arctan\left(\frac{y_1}{y_2}\right)\cdot L}{2\pi} + L, L\right), \tag{1}$$

where L is the height of the simulation box and the mod functions mod(A,B) returns the remainder of A when divided by B.

$$y_1 = \sum_{i=1}^{N} \sin\left(\frac{2\pi}{L}\cdot z_i\right), \tag{2}$$

$$y_2 = \sum_{i=1}^{N} \cos\left(\frac{2\pi}{L}\cdot z_i\right). \tag{3}$$

Additionally, we analyzed the equilibration through the concentration profiles. We divided the simulation trajectories into 10 intervals. Figure SI 2 shows the concentration profiles converge within 30 percent of the simulations for both initial conditions.

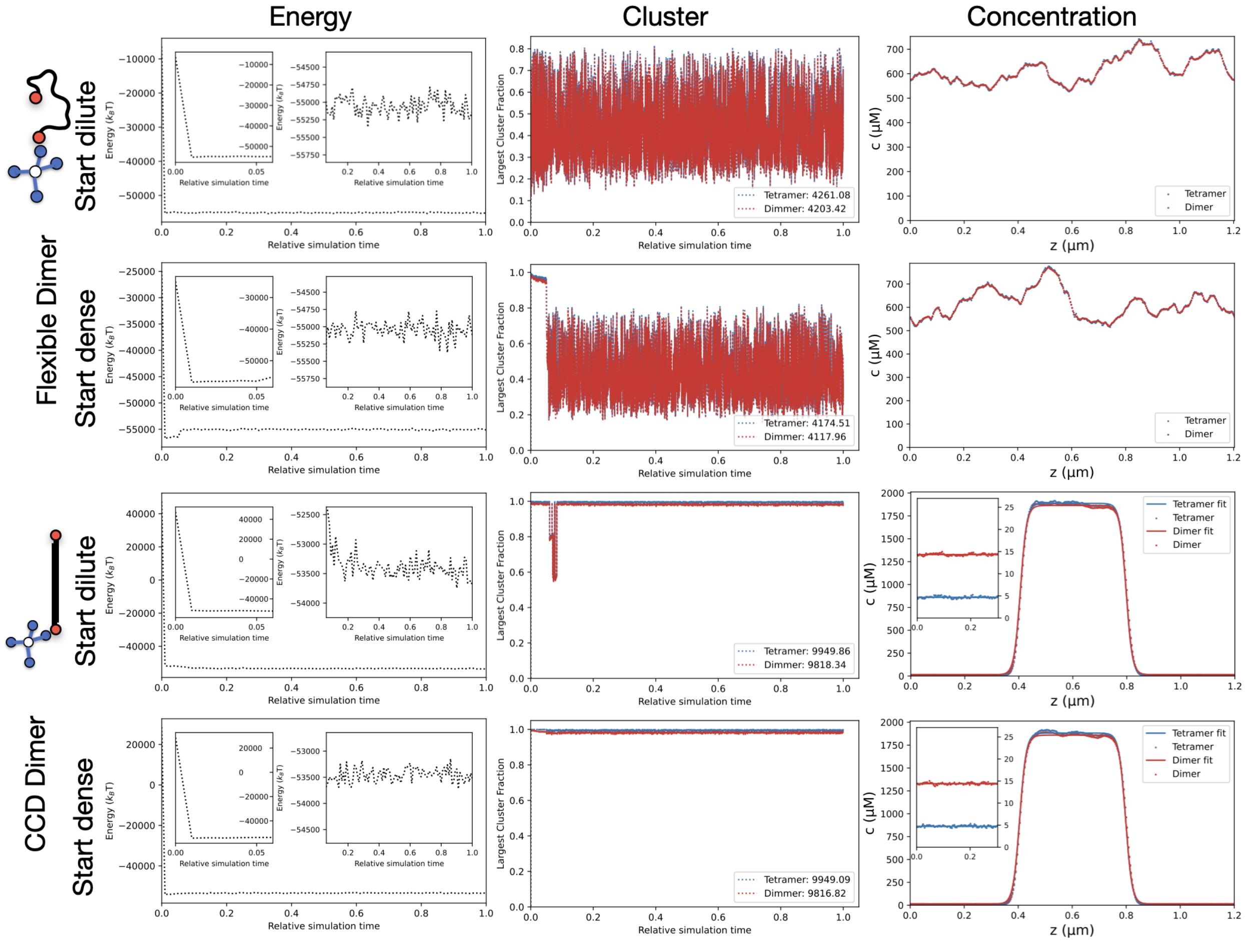


Figure 1: **Convergence of simulations starting from different conditions.** Columns (left to right) show (i) total energy of the system, (ii) largest cluster size as simulation continues, which shows the system equilibrates within the first 10% of the simulation time; and (iii) final molecule distributions along z-axis. Both dilute and dense starting conditions lead to the same thermodynamic equilibrium state. The elongation potential determines the phase behavior

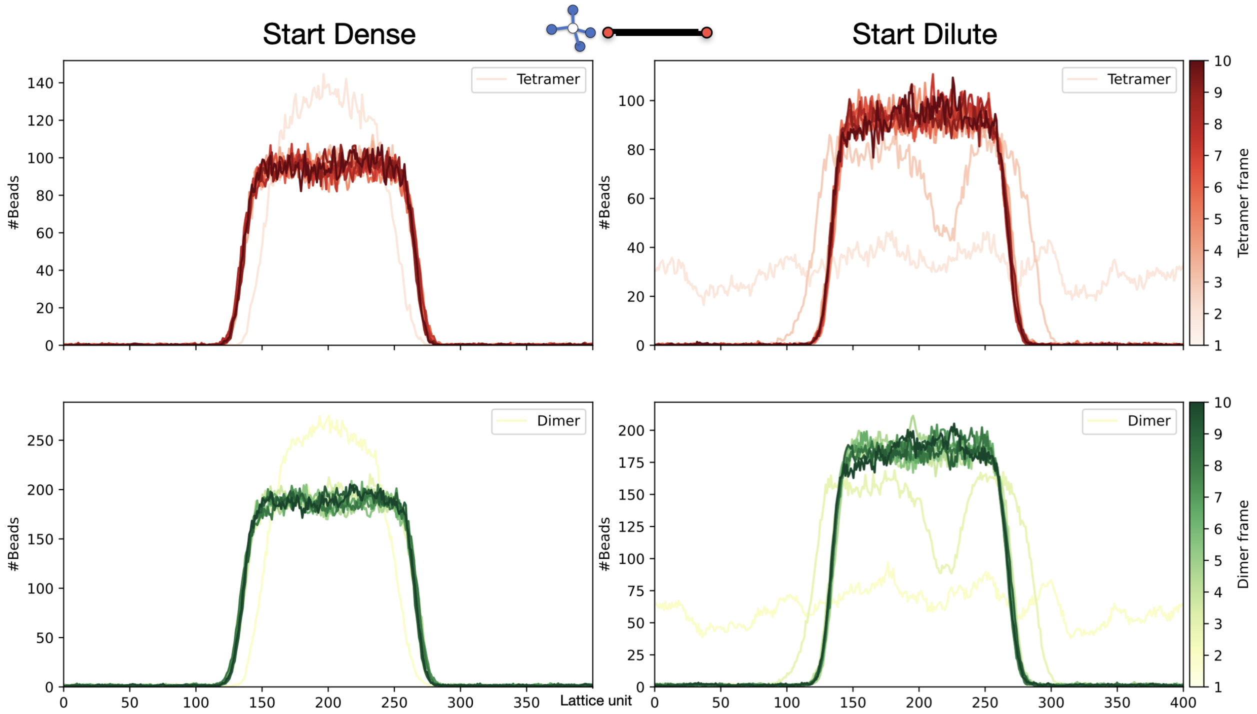


Figure 2: **Time-resolved convergence of oligomer distributions along z-axis.** The full simulation is divided into 10 equally spaced time intervals, with the oligomer distribution for each interval averaged and shown as one frame. Intervals closer to the final frames are represented with more intense colors. Simulations starting from both dilute and dense conditions converge to the same equilibrium distribution within the first half of the simulation.

### Determining gelation transition and liquid-liquid phase separation

Three phases are possible for polymer systems: a mixed solution (sol), a gel state without LLPS (gel), or a gel state with LLPS (condensate).

A gel is defined as a cross-linked polymer network spanning the entire volume of the system, which corresponds to the formation of an infinitely large cluster in an infinite system. Similar to the meaning of $c_{\mathrm{sat}}$, gel point concentration ($c_{\mathrm{gel}}$) is the minimum required for the system to form a gel.

We used the concentration profiles along z-axis as the primary metric for gelation versus LLPS. Additionally, this is complemented by cluster size analysis.

We note, for systems that undergo gelation but not LLPS, the 're-center' procedure is not suitable anymore. This is because it causes an artifact of increased density in the middle of the simulation box due to accumulation of statistical noise (Figure SI 3b and c).

#### Phase separation

Phase separation was assessed quantitatively by extracting the dense phase concentration and the dilute phase concentration from the concentration profile along z-axis:

1. We fit the concentration profiles using a symmetric sigmoid function, which is defined as $f(z) = -\frac{a}{1+e^{b\cdot(z-(L/2-l))}} + \frac{a}{1+e^{b\cdot(z-(L/2+l))}} + d$, where L is the length of simulation box in z-dimension.

2. We extracted from the fit $c_{\mathrm{dense}} = f(L/2)$ and $c_{\mathrm{dilute}} = f(L)$.

3. Phase separation was identified when the partition factor, $P = c_{\mathrm{dense}}/c_{\mathrm{dilute}}$, is greater than 3. When $P$ is smaller than 3, the system may or may not undergo gelation (see next section).

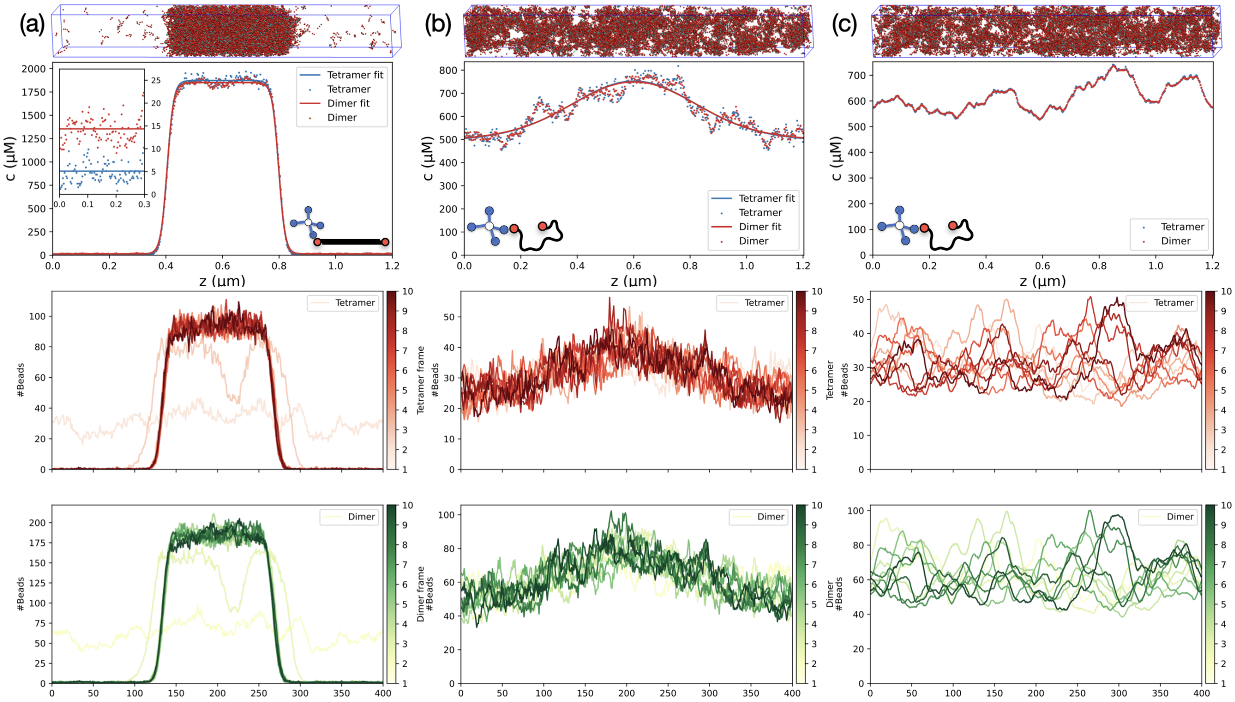


Figure 3: **Liquid liquid phase separation or Gelation** Top: Snapshot of a SLAB simulation and the average concentration profile along z-axis. Middle and Bottom: Time-resolved oligomer distribution of tetramer and dimer. The whole simulation is equally separated by 10 period, and the distribution of each period is averaged as one frame. (a) CCD Dimers. The concentration profile shows clear dense and dilute phases, confirming liquid-liquid phase separation (LLPS). (b) Flexible Dimers with 're-center'. The profile shows no phase separation, with a slight artifact due to translational alignment. (c) Flexible Dimers without 're-center'. The distribution remains homogeneous without signs of phase separation.

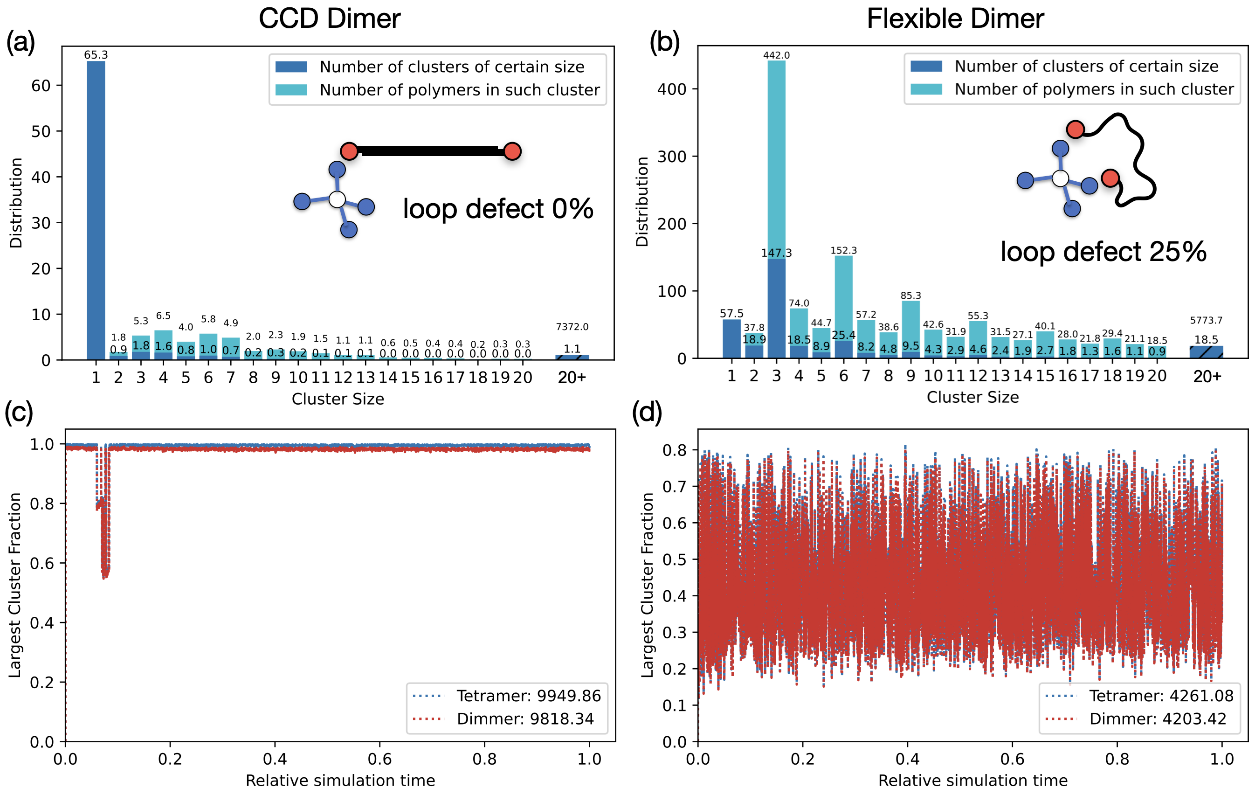


Figure 4: **Cluster size distribution and largest cluster size fraction** There are 2500 tetramers and 5000 dimers in a box size of 50*50*400 lattice unit. (a) Cluster size distribution for CCD Dimers, meaning the number of clusters of certain size. There are also 65 polymers that are not connected in a cluster, and some other small clusters. The last column stands for the number of clusters that contain more than 20 polymers, which is averagely 1. Namely, this the condensate, which contains 7372 polymers. The fraction of loop defect (tetramers in a closed state) is also calculated, which is near 0%. (b) Cluster size distribution for Flexible Dimers, showing a broader distribution with many clusters of different sizes. The most populated cluster size 3 corresponds to the closed state where one tetramer is fully closed by two dimers. The last column indicates that there are about 18 clusters that contain more than 20 polymers. Those condensates intotal contain 5774 polymers. The fraction of loop defect is around 25%. (c) Fraction of oligomers in the largest cluster for CCD Dimers. The value one indicates that almost all molecules are in the largest cluster (condensate). The flat behavior indicates the stable clustering. (d) Fraction of oligomers in the largest cluster for Flexible Dimers, showing significant fluctuations, as a characteristic of gelation.

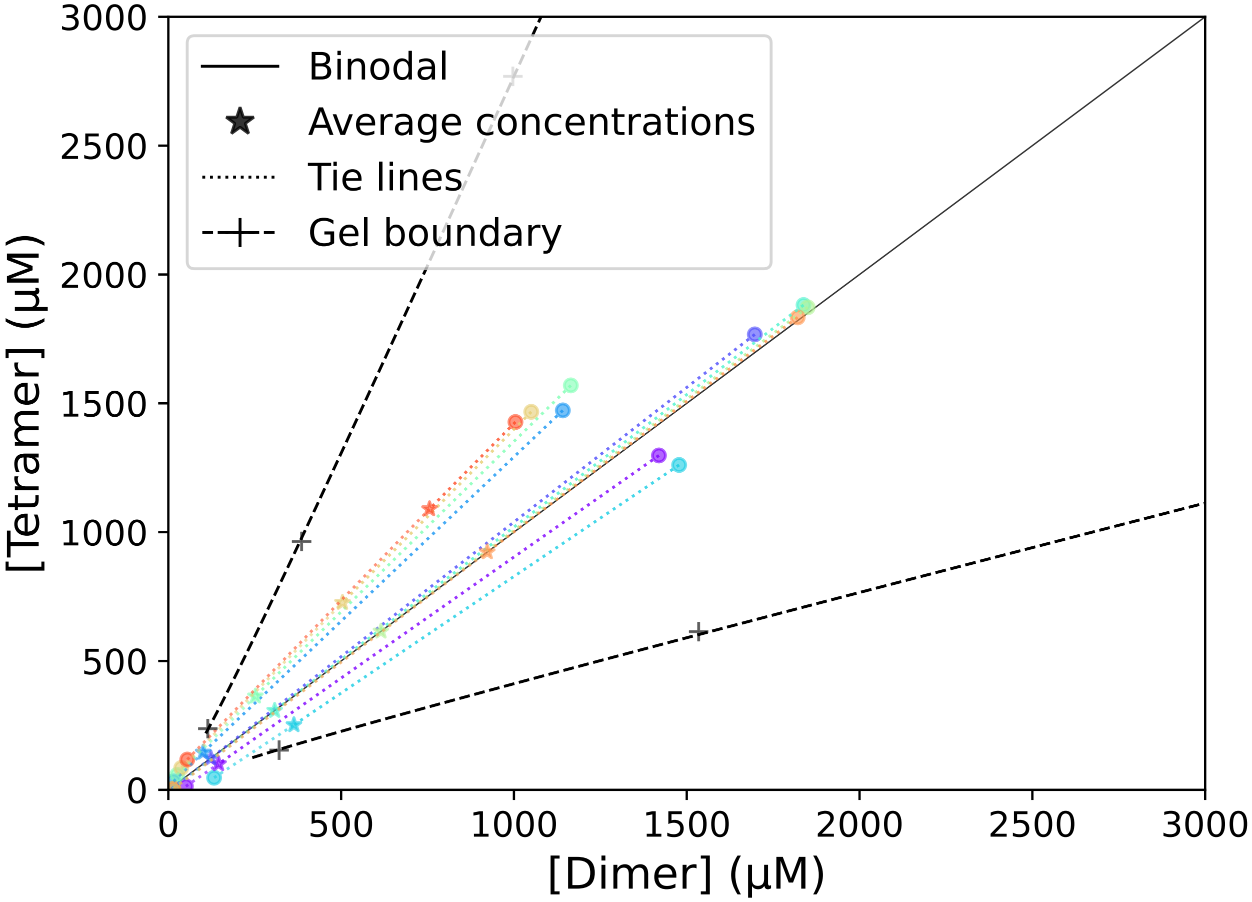


Figure 5: **Phase diagram in linear scale (Figure 4b in main text)** Phase diagram for CCD Dimers at equilibrium length Leq = 21 nm. Simulations with average concentrations in the demixing region robustly phase separate. Each average concentration and its corresponding dense and dilute phase points are connected by a color-matched tie line (dotted line). The binodal (phase boundary) was determined by the resulting concentrations of the two coexisting phases in the demixing region. Simulations in the gel region form a gel without phase separating.

#### Gelation

We define gelation as when the largest cluster exceeds a critical cluster size threshold motivated by Flory-Stockmayer gel theory.[2,3]

Flory-Stockmayer theory predicts the percolation threshold based on component valences and binding probabilities for an infinite size system. The critical parameter governing gelation is the average number of additional components recruited into a growing network, denoted as $\epsilon$. For a single component with valence $V_n$ and occupancy fraction $x$, this is given by $\epsilon = (V_n - 1)x$.

For a binary system (e.g., our tetramer-dimer system) with components A and B of valences $V_{nA}$ and $V_{nB}$, the overall $\epsilon$ is:

$$\epsilon = \epsilon_A \epsilon_B = (V_{nA} - 1)x_A(V_{nB} - 1)x_B. \tag{4}$$

where $x_A$ and $x_B$ represent the fraction of occupied binding sites for A and B, respectively. Gelation occurs when $\epsilon > 1$, indicating that, on average, every new component incorporated into the network recruits more than one additional component, leading to a cascade of network expansion. Conversely, $\epsilon < 1$ results in network termination and no gelation.

The fraction of bound sites can expressed as:

$$x_A = \frac{[AB]}{[A]}, x_B = \frac{[AB]}{[B]}, \tag{5}$$

where [AB] is the concentration of bound complexes between A and B. The equilibrium dissociation constant is given by:

$$K_d = \frac{([A] - [AB])([B] - [AB])}{[AB]} \tag{6}$$

$$K_d = \left(\frac{[A]}{[AB]} - 1\right)\left(\frac{[B]}{[AB]} - 1\right)[AB] \tag{7}$$

The onset of the gelation transition occurs when

$$\epsilon = 1 = (V_{nA} - 1)x_A(V_{nB} - 1)x_B = \lambda x_A x_B, \tag{8}$$

where $\lambda = (V_{nA} - 1)(V_{nB} - 1)$ defines the product of effective valence contributions. Substituting equation 5 into equation 8 we have

$$[AB] = \sqrt{\frac{[A][B]}{\lambda}}. \tag{9}$$

By substituting this into equation 7, we have:

$$K_d = \left([A]\sqrt{\frac{\lambda}{[A][B]}} - 1\right)\left([B]\sqrt{\frac{\lambda}{[A][B]}} - 1\right)\sqrt{\frac{[A][B]}{\lambda}}. \tag{10}$$

This can be rewritten as

$$K_d = \left(\sqrt{\frac{\lambda}{r}} - 1\right)(\sqrt{r\lambda} - 1)\sqrt{\frac{r}{\lambda}}[A], \tag{11}$$

where $r = \frac{[B]}{[A]}$ is the ratio of two components. Then the gel point of component A is

$$[A] = \frac{K_d\sqrt{\lambda}}{(\sqrt{\lambda} - \sqrt{r})(\sqrt{r\lambda} - 1)}. \tag{12}$$

For a 1:1 stoichiometric system (r = 1), the gel point concentration simplifies to:

$$[A] = [B] = \frac{K_d\sqrt{\lambda}}{(\sqrt{\lambda} - 1)^2}. \tag{13}$$

We used this gel point concentration from equation 9 to calculate the size of a cluster in our finite-size simulation that corresponds to a infinite gel: we randomly assigned $N_{AB} = \sqrt{\frac{N_A N_B}{\lambda}}$ bonds, where $N_A$ and $N_B$ are the numbers of interaction domains. Adding the polymer connectivity, we found the number of polymers in the largest connected network. This was averaged over 10,000 independent realizations, giving the critical cluster size for a percolated gel.

We note this threshold does not include the linker/polymer properties and the slab geometry. We expect this is a good estimate for finite size corrections. The resulting percolation thresholds are shown in Table SI 1.

Table 1: **top10 cluster size at gel point** According to Flory-Stockmayer theories,[2–4] there is a threshold of cluster size for the system to undergo gel transition. The top 10 large cluster size, the gel threshold, at different ratio for total of 20000 binding sites are given here.

| **Ratio** | **1** | **2** | **3** | **4** | **5** | **6** | **7** | **8** | **9** | **10** |
|---|---|---|---|---|---|---|---|---|---|---|
| 0.36 | **603.03** | 263.32 | 177.59 | 137.07 | 113.19 | 97.18 | 85.76 | 77.07 | 70.22 | 64.65 |
| 0.40 | **607.44** | 265.62 | 178.91 | 138.15 | 114.11 | 98.10 | 86.52 | 77.76 | 70.84 | 65.22 |
| 0.48 | **612.97** | 268.68 | 181.01 | 139.88 | 115.59 | 99.36 | 87.68 | 78.80 | 71.80 | 66.11 |
| 0.69 | **622.63** | 272.51 | 183.68 | 142.03 | 117.39 | 100.94 | 89.08 | 80.09 | 72.99 | 67.22 |
| 1.00 | **627.38** | 273.74 | 184.12 | 142.32 | 117.62 | 101.14 | 89.27 | 80.25 | 73.14 | 67.37 |
| 1.44 | **621.35** | 270.00 | 181.76 | 140.46 | 116.09 | 99.84 | 88.11 | 79.22 | 72.21 | 66.52 |
| 2.08 | **606.62** | 262.85 | 176.75 | 136.60 | 112.91 | 97.11 | 85.71 | 77.08 | 70.28 | 64.75 |
| 2.50 | **591.70** | 256.95 | 173.14 | 133.84 | 110.72 | 95.25 | 84.10 | 75.63 | 68.96 | 63.53 |
| 2.78 | **589.04** | 254.95 | 171.32 | 132.36 | 109.38 | 94.10 | 83.12 | 74.72 | 68.11 | 62.74 |

We used these cluster size thresholds to calculate the gel points in our simulations by adjusting the height of the simulation box to find the concentration when the observed cluster size matches the threshold.

The gel lines shown in Figure 4 from the main text were calculated for different tetramer: dimer binding site ratios. Panels (a) and (c) in Figure SI 6 show data used to calculate the threshold concentrations for CCD Dimer and Flexible Dimer systems respectively. Panels (b) and (d) in Figure SI 6 show the interpolation of the results to find the gel lines using the function form

$$c_{\mathrm{gel}}(r) = c\left(\frac{a}{(\log r + \log 3)^{\lambda_1}} + \frac{a}{(\log 3 - \log r)^{\lambda_2}} - \frac{a}{(\log 3)^{\lambda_1}} - \frac{a}{(\log 3)^{\lambda_2}} + 1\right). \tag{14}$$

These fit functions were used to plot the gel line in the phase diagrams (Figure 4b and c in main text).

There are two singular points in the $c_{\mathrm{gel}}$, which are when $r = 3$ and $r = 1/3$. This means when $r$ is approaching 3 or $1/3$, the system needs be infinite dense to form a gel (a cluster whose size exceeds the Flory-Stockmayer threshold). This is because the ratio of effective valency for network recruitment is (4-1):(2-1)=3:1. This defines the stoichiometry ratio window [1/3,3], only within which the system can form a gel.

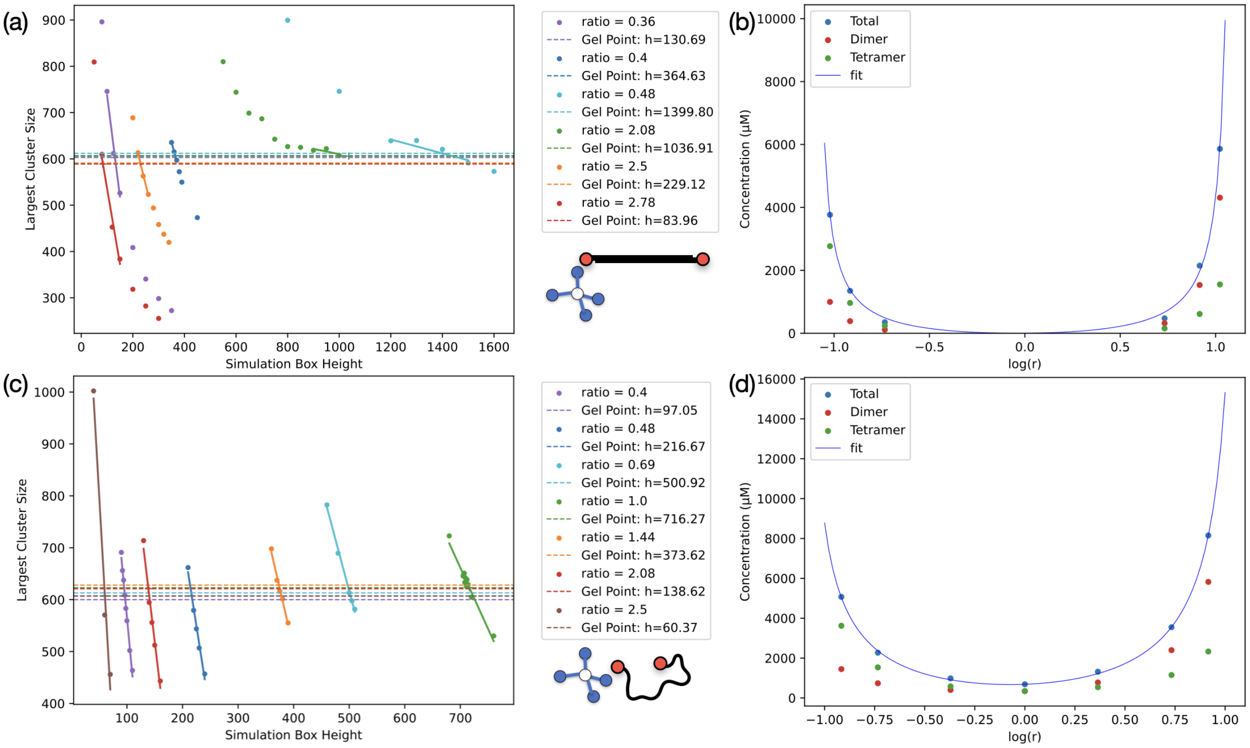


Figure 6: **Determination of gel line by gel points at varying [Dimer]:[Tetramer] ratios $r$ and simulation box** (a) and (c) Largest cluster size as a function of simulation box height for CCD Dimers and Flexible Dimers, respectively. Gel points were identified where the largest cluster size equals the critical cluster size ( $CS^*$ ) from Flory-Stockmayer theory. Each system contains 20000 binding sites in total, where the ratio is adjustable. Simulation box height, which determines the concentration, is the parameter tuned for each ratio to get the gel point. (b) and (d) Concentration of gel points and the corresponding fit to the functional form $c_{\mathrm{gel}}(r)$.

### Impact of elongation potential on phase behavior

The elongation potential plays a crucial role in modulating phase behavior. We simulated systems with varying CCD equilibrium lengths $L_{eq}$. There is a critical threshold $L^*_{eq} \approx$ 12 nm separating phase behavior-'LLPS' or 'gelation without LLPS'. For each $L_{eq}$, the corresponding representative phase diagrams are plotted based on the functional form of $c_{\text{gel}}(r)$ as a guide to the eye (Figure SI 7b). As $L_{eq}$ increases above $L^*_{eq}$, the binodal shifts to lower concentrations . This indicates that longer CCD lengths promotes LLPS. Inversely, as $L_{eq}$ decreases below $L^*_{eq}$, the gel line shifts to higher concentrations. This indicates that shorter CCD lengths weaken gelation, as the gel point concentration increases.

To further explore the effect of the elongation potential, we investigated how varying the strength of the elongation potential coefficient $K$ influences phase behavior. As the elongation constraint becomes more strict ($K$ increases), the geometric frustration preventing loop defects becomes stronger (Figure SI 7 c). The effect plateaus when $K \geq 1 k_B T/LS^2$. As long as $K$ is an appreciable strength, then the phase behavior is primarily governed by the $L_{eq}$ (Figure SI 7 d).

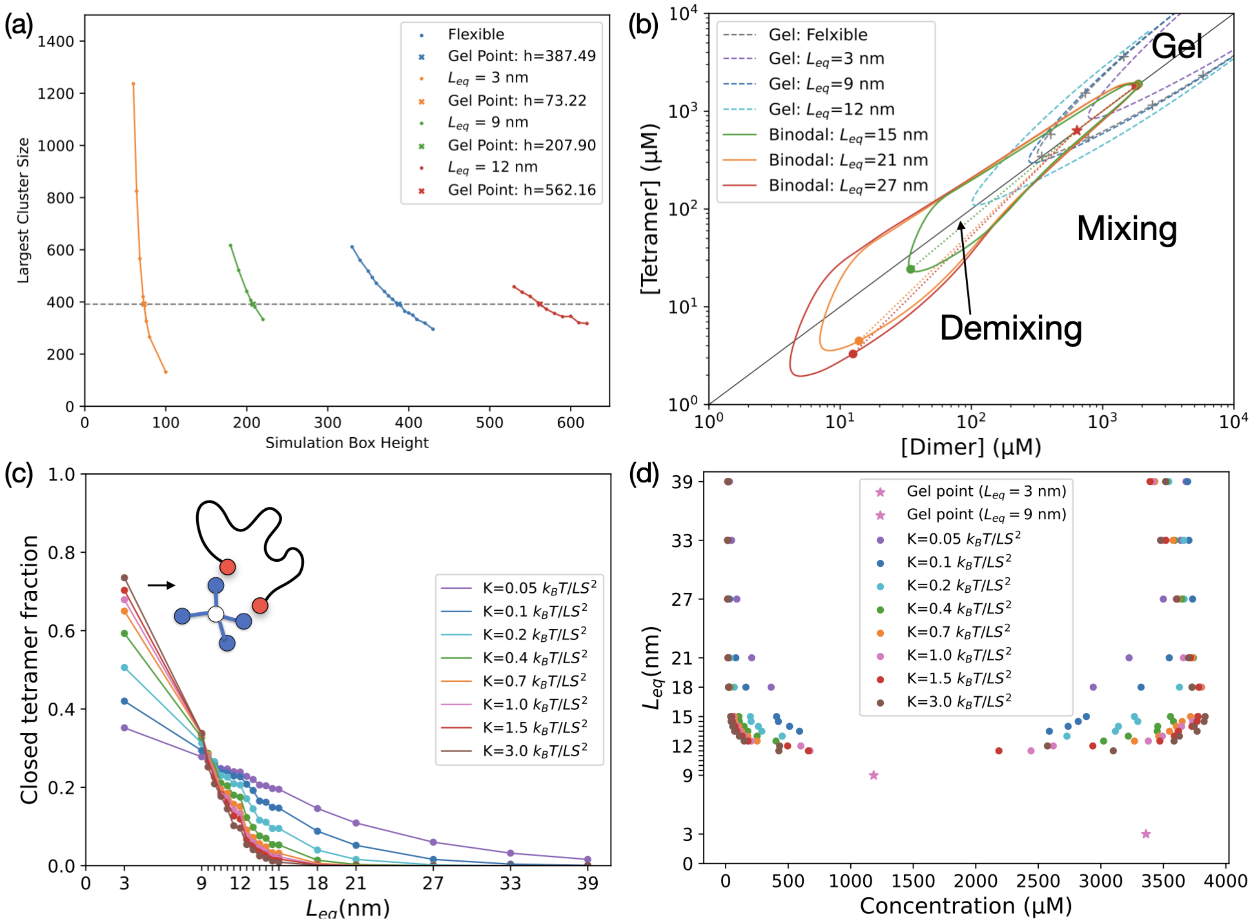


Figure 7: **Phase behavior at different elongation potential** All simulations are at 1:1 stoichiometry. (a)Gel points at varying CCD equilibrium length $L_{eq}$ with elongation potential coefficient $K = 1k_BT/LS^2$. Shorter $L_{eq}$ values increase the gel point concentration, weakening gelation. (b) Phase diagrams for different $L_{eq}$ values. Systems with $L_{eq} < 12$ nm undergo gelation without LLPS (dashed lines). Systems with $L_{eq} > 15$ nm phase separates (solid lines). The shape of gel lines and binodal are based on and rescaled from Figure 4 to match the results from simulations at 1:1 stoichiometry with varying $L_{eq}$. These act as a guide to how the phase diagrams are expected to behave as a function of CCD length. (c) Fraction of closed tetramers as a function of $K$. Results show that $K$ has no qualitative impact on tetramer closure. (d) Phase diagram showing the relationship between $L_{eq}$, total component concentrations (dilute and dense), with varying elongation potential strengths $K$.

#### Conversion of binding energy to $K_d$

To calibrate the relationship between $K_d$ and $\epsilon$, we simulated the equilibrium of E9 and Im2 interactions as monomeric beads with varying $\epsilon$ from $-2k_BT$ to $-14k_BT$. Here $K_d$ is the dissociation constant, defined as

$$K_d = \frac{[E9][Im2]}{[E9:Im2]} \tag{15}$$

where [X] denotes the concentration of species X.

The relation of $K_d$ and binding energy $\epsilon$ follows

$$K_d \approx Ae^{\epsilon/k_BT} \tag{16}$$

where $A$ is constant.

Figure SI 8 shows the linear fit of $\log(K_d)$ as function of $\epsilon$, yielding $A = -6.042$.

Notably, the wild-type (WT) binding affinity of the E9-Im2 interaction corresponds to a binding energy of:

$$\epsilon^{WT} = \log(1.5 \times 10^{-8}) + 6.042 = -11.973\, k_BT. \tag{17}$$

This binding energy is too strong for efficient simulations. On average, breaking a single bond at this energy scale would require $\approx e^{11.973} = 157,945$ moves, which is computationally impractical.

To address this, we chose a binding energy of $\epsilon = -6\, k_BT$ for simulations, corresponding to a binding affinity of:

$$K_D(\epsilon = -6\, k_BT) = e^{-12.042}\, \mathrm{M} = 5.89 \times 10^{-6}\, \mathrm{M}. \tag{18}$$

In thoery, this choice introduces a concentration discrepancy of:

$$\frac{C_{\text{simu}}}{C_{\text{WT}}} \approx e^{-6-(-11.973)} = e^{5.973} = 392.68 = 10^{2.59}. \tag{19}$$

Namely, our simulations yield concentrations that are 2.59 orders of magnitude higher than those in experiments. Comparing the phase diagram of experimental measurements and our simulations (Figure 4a and b), one can observe roughly a discrepancy of 3 orders of magnitude. This suggests that our simulations are in remarkably quantitative agreement with experimental measurements, especially considering the minimalistic nature of the simulations, zero fitting parameters, and the numerous factors not explicitly accounted for.

Table 2: **Reported Binding Affinity of E9-Im2 Variants.**
Previously reported[5] mean and standard error values of the affinities are given ($n = 2$), as used in work by Heidenreich et al.[6]

| **Im2 Mutation** | $K_d$ **with E9 (M)** |
|---|---|
| **D33L N34V R38T** | $3.4 \pm 1.4 \times 10^{-13}$ |
| **D33L** | $4.8 \pm 0.3 \times 10^{-11}$ |
| **N34V R38T** | $1.9 \pm 0.4 \times 10^{-10}$ |
| **R38T** | $2.6 \pm 0.5 \times 10^{-9}$ |
| **N34V** | $3.3 \pm 0.7 \times 10^{-9}$ |
| **WT** | $1.5 \pm 0.1 \times 10^{-8}$ |
| **E30A** | $2.8 \pm 1.6 \times 10^{-7}$ |
| **P56A** | $2.1 \pm 0.7 \times 10^{-6}$ |
| **V37A** | $9.3 \pm 4.4 \times 10^{-6}$ |

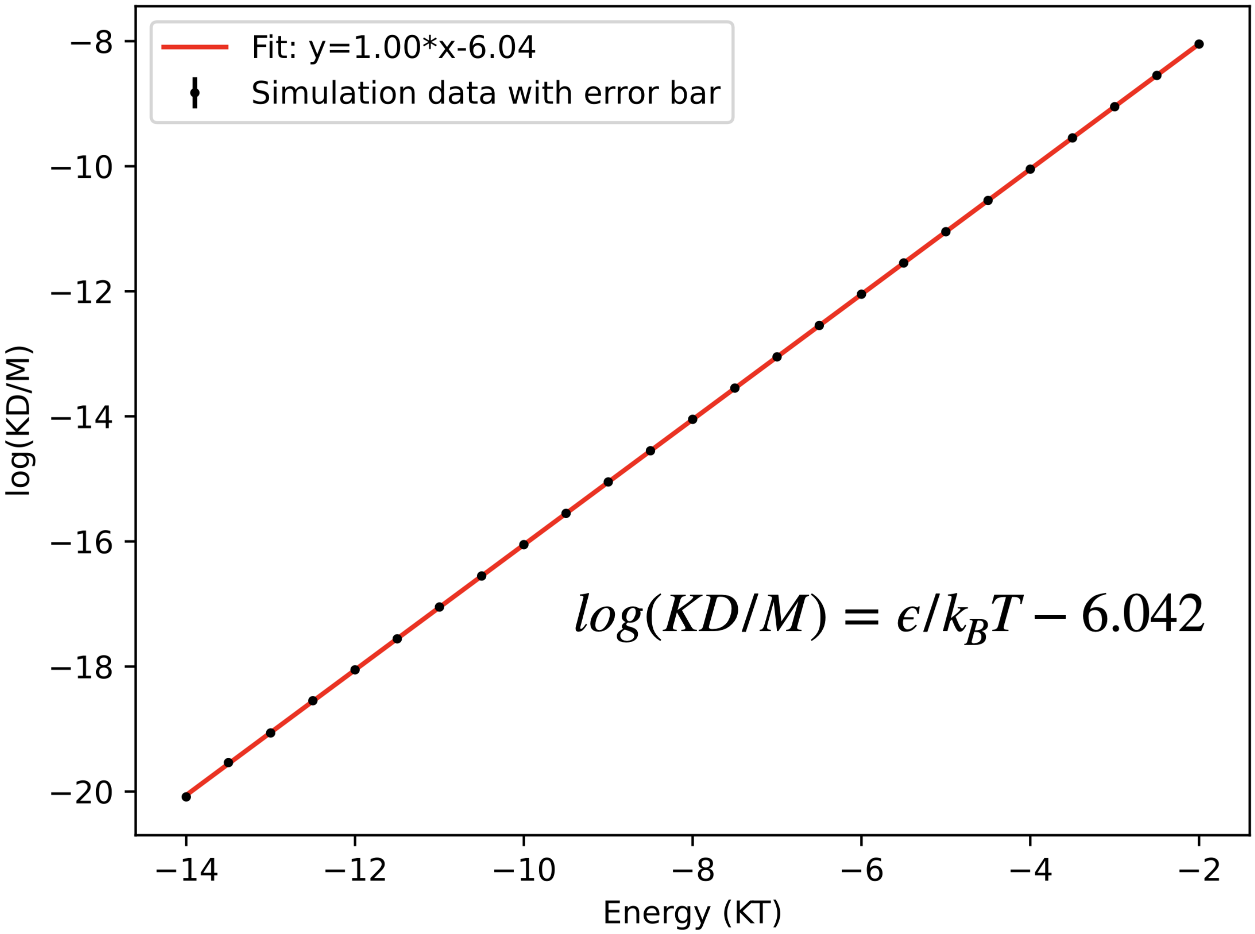


Figure 8: **Binding Strength ($K_d$) vs. Binding Energy Used in Simulations.** Simulations were conducted with monomeric E9 and Im2 interacting beads to determine the relationship between binding energy ($\epsilon$) and dissociation constant ($K_d$).
